# Assessment of Ex Situ Conservation Strategies for the Preservation of Domestic South American Camels in Southern Peru: Insights from Mitochondrial DNA Analysis

**DOI:** 10.1101/2025.06.26.661050

**Authors:** Orson Mestanza, Diogenes Cerna, Claudia E. Yalta-Macedo, Teodosio Huanca, Norberto Apaza, Evelyn Díaz Salas, Adriana Vallejo-Trujillo

## Abstract

Southern Peru is home to the largest population of alpacas and the second-largest population of llamas. For many rural families, breeding these animals is vital. Recently, breeding communities have faced several challenges, including restricted market access, climate change, a reduction in coloured individuals, and pasture degradation, which hinder trading and genetic improvement efforts. This study examines the D-loop region of mitochondrial DNA to assess the genetic maternal diversity of alpacas and llamas. It also evaluates this alongside the preservation initiatives of the Quimsachata ex situ germplasm bank, which aims to sustain both phenotypic and genetic diversity while advancing breeding technologies. Our analysis identified 55 maternal haplotypes with 41 variation sites, indicating significant genetic diversity (*h* ≈ 0.8–0.9) and low differentiation (*Fst* = 0.01–0.05) within these populations. This finding is particularly noteworthy when compared to the maternal haplotypes of domestic and wild South American camelids (SAC) in other areas of South America. We did not observe any recent demographic shifts in population structure. Additionally, our research uncovers two genetic maternal clusters among alpacas and llamas, suggesting their ancestral connections to vicunas and guanacos, with indications of crossbreeding, particularly among alpacas, which show a closer lineage to guanaco-llamas. The Quimsachata bank currently safeguards a significant diversity of maternal heritage in alpacas and llamas within this region. The monitoring and protection of these resources is crucial for the economic advancement of Andean communities and can help promote genetic programmes aimed at enhancing sustainable productivity.

## INTRODUCTION

Domestic camels of South America, specifically alpacas (*Vicugna pacos*) and llamas (*Lama glama*), have served as vital herding animals for communities in the Andean highlands for 5,000 to 6,000 years since their domestication (Yacobaccio, 2021). During the Inca Empire, these animals were essential sources of wool and meat (Gade, 1996) and played a crucial role in transportation and trade, thriving in the harsh Andean terrain (Wheeler *et al*., 1995). Today, they remain indispensable to Andean populations. Alpacas support the livelihoods of approximately 82,000 rural families living in extreme poverty in the Peruvian Andes, while llamas assist about 95,000 families (Wurzinger & Gutierrez, 2022).

Alpacas and llamas produce highly valued fibre for the textile industry, with exports exceeding US$14 million in 2023 (PROMPERÚ, 2023). They also provide healthy, low-cholesterol meat, and their hides are utilised in the tanning industry (MINAGRI, 2012; MIDAGRI, 2024). Additionally, llama manure serves as both fuel and fertiliser, making these animals essential as pack animals in high-Andean cultures and for tourism (Fundación WIESE, 2018).

South America is home to over 12 million camelids, including 4 million llamas and 7.5 million alpacas. Peru boasts the world’s largest alpaca population, at 49%, and ranks second in llamas, at 19% (FAO, 2024; INEI, 2012). The high-quality fibre of alpacas and llamas is available in over 22 natural colours. However, the industry’s demand for white fibre has led to intense selection processes, resulting in a loss of genetic diversity, particularly due to the decrease in coloured (non-white) individuals as the focus has shifted towards increasing the number of white alpacas and llamas (Anello *et al*., 2022; Fernandez-Baca, 1994).

In response to growing concerns regarding the whitening of alpaca herds amongst small, medium, and large breeders, the National Institute of Agrarian Innovation (INIA) of the Peruvian government established an ex situ germplasm bank in Illpa, Puno Region, in 1988 through its Camelid Research Programme. This initiative primarily aims to recover coloured Suri and Huacaya alpacas, as well as llamas in their two ecotypes: Chaku and K’ara (Huanca *et al*., 2007). This project represents the largest ex situ conservation effort for South American camels (SAC), significantly enhancing the preservation of their genetic diversity. By collaborating with smallholders, it effectively supports in situ conservation of both genetic and phenotypic diversity (Paredes *et al*., 2020; Huanca *et al*., 2007).

Numerous studies have examined genetic diversity and evolution, linking llamas to guanacos (*Lama guanicoe*) and alpacas to vicunas (*Vicugna vicugna*) (Diaz-Maroto *et al*., 2021; Fan *et al*., 2020; Sarno *et al*., 2001; Marin *et al*., 2007, 2008). Notably, Wheeler *et al*. (1995) provided evidence of post-conquest hybridisation between llamas and alpacas through fibre analysis. Moreover, research on mitochondrial control region sequences across South America indicated interbreeding during domestication by early herders (Diaz-Maroto *et al*., 2021). Hybridisation between these species has also been observed in alpacas and llamas from Bolivia (Echalar & Barreta, 2022) and Polish alpaca populations (Podbielska & Piorkowska, 2022).

The southern region of Peru, home to the Quimsachata germplasm bank, is the site of approximately 50% of the country’s alpacas and llamas (INEI, 2012). Paredes *et al*., (2020) reported a high level of genetic diversity within the Quimsachata llama population. However, there is a lack of research on ex situ alpaca populations and a need for further studies on populations throughout the southern region, particularly since this area has the highest density of these species and provides most of the breeding animals introduced to other regions of the country (FAO, 2005).

In this study, we analyse the D-loop mitochondrial DNA (mtDNA) of alpacas and llamas from the Quimsachata ex situ germplasm bank, as well as from herds belonging to small and medium-sized holders across the two main agroecological zones (dry and wet puna) in southern Peru. Our objectives are to: (i) evaluate the maternal haplotype diversity present within herds, (ii) assess the extent of hybridisation between the two species in the region, (iii) evaluate the role of the germplasm bank in preserving the genetic diversity of these species, and (iv) contribute to decision-making regarding conservation efforts aimed at improving sustainable production programmes at both regional and national levels.

## METHODOLOGY

### Sampling

Blood and hair samples, along with a photographic record, were collected from a total of 407 alpacas, representing 51 herds, and 282 llamas from 33 herds (refer to Supplementary Table S1 for details). These animals were sourced from various locations across the Puno region, as well as from regions of Cusco, Tacna, and Moquegua, which are situated in the southern highlands of Peru. The sample collection comprised both Suri and Huacaya alpacas (*Vicugna pacos*) and Chaku, K’ara, and Suri llamas (*Lama glama*).

The samples were classified according to the agroecological zones to which the animals belonged: the wet Puna in northern Puno, characterised by an extended wet season of eight months, higher rainfall (ranging from 400 to 700 mm per year), and the presence of montane grasslands and shrublands; and the dry Puna in southern Puno, which has a shorter wet season of two months, lower rainfall (less than 400 mm per year), and high-altitude grasslands (Oyague & Cooper, 2020; Huanca *et al*., 2007) (see Figure 1). Additionally, samples from 39 llamas and 44 alpacas were obtained from the “Quimsachata” Germplasm Bank at the Illpa Experimental Station of the National Institute of Agrarian Innovation (INIA).

**Figure 1.**
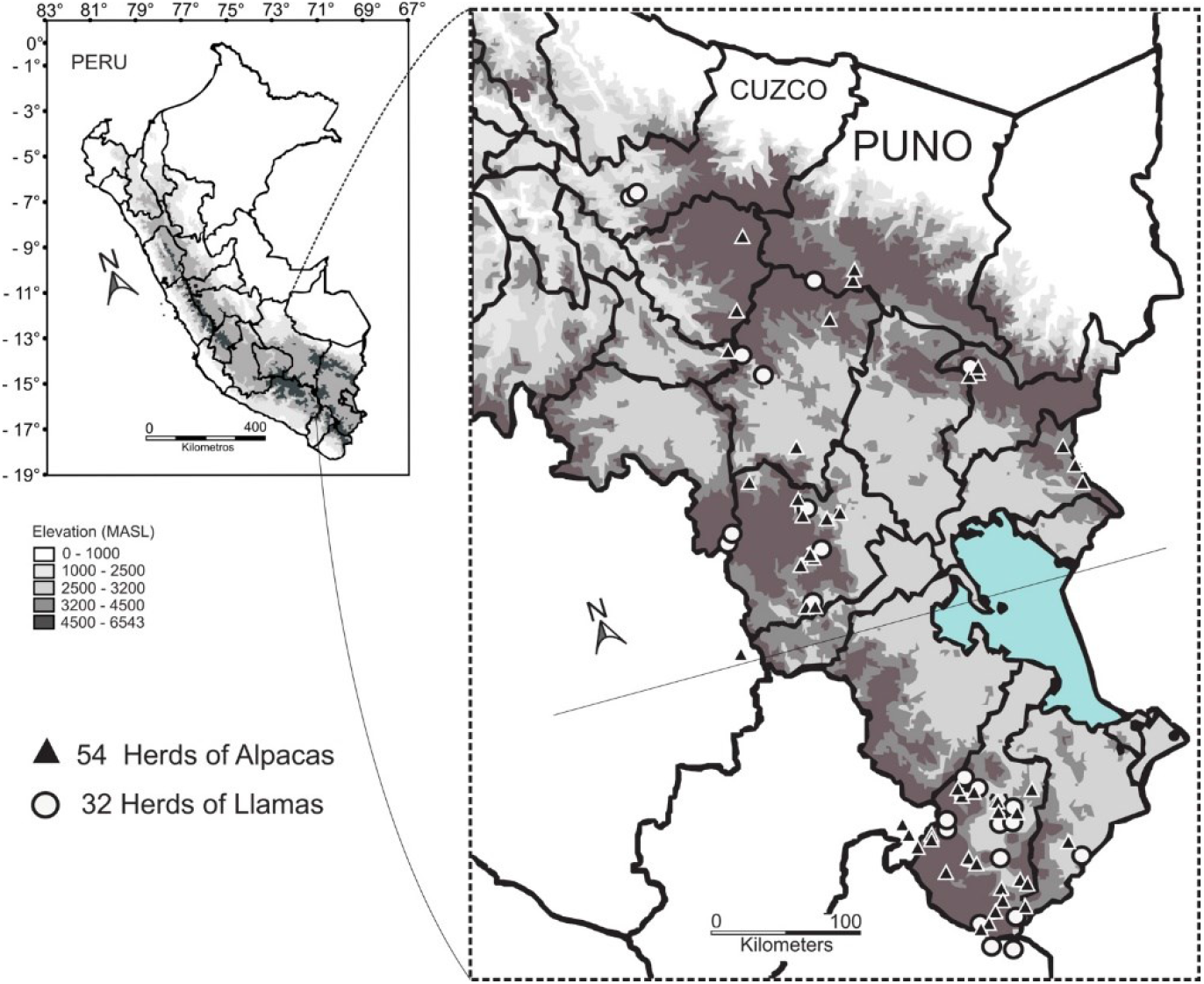
Geographic origin of collected samples from the Southern region of Peru (Puno, Cusco, Tacna, and Moquegua departments)

### DNA Extraction and Sequencing

DNA was extracted using a modified version of the chloroform-isoamyl alcohol protocol for hair (Sambrook *et al*., 1989), which included the addition of Dithiothreitol (DTT). Blood samples were collected on FTA cards (Whatman Bioscience), and DNA extraction was performed according to established laboratory protocols that utilised 5% Chelex resin.

The hypervariable segment I of the D-loop region, spanning 513-514 bp, was amplified using the forward primer LThrArtio (5’-GGTCTTGTAAAACGAAGCAGGA-3’) and the reverse primer H15998 (5’-CGCAGTCAAATCAATTGAAGCTGG-3’) described by Marin *et al*., 2007. Amplification was performed in a final volume of 25 µL, containing 30–40 ng of DNA. The thermal cycling profile comprised an initial denaturation at 94 °C for 5 minutes, followed by 35 cycles of denaturation at 94 °C for 45 seconds, annealing at 64 °C for 45 seconds, and extension at 72 °C for 1 minute. A final extension step was carried out at 72 °C for 11 minutes. The PCR products were purified and sequenced using an automated ABI 377 sequencer (Applied Biosystems), employing the same primers that were used for amplification. The resulting sequences were deposited in GenBank under project 2939020 (accession numbers PV359989-PV360677).

### Genetic diversity and population differentiation analyses

A total of 688 sequences were assembled using SangeranalyseR v.1.14 (Chao *et al*., 2021). Ambiguous positions were curated with SeqScape v2.5 (Applied Biosystems), utilising the reference sequences from GenBank NC_012102.1 for llamas and EU285663.1 for alpacas. The sequences were aligned, and extreme values with gaps were discarded. This alignment was then used to calculate genetic diversity indices, including the number of variable sites, the number of haplotypes, haplotype diversity, nucleotide diversity, and transversion among populations. These calculations were performed using Pegas v.1.3 (Paradis, 2010).

To understand the demographic history of domestic camelids, we performed neutrality tests, including Tajima’s D (Tajima, 1989) and R2 tests (Ramos-Onsins & Rozas, 2002). We also calculated the genetic distance between populations using Nei’s and Fst population distance metrics, utilising the adegenet v2.1.10 (Jombart, 2008) and hierfstat v.0.5.11 (Goudet, 2005) packages, respectively. Additionally, we computed the observed mismatch distribution of pairwise distances.

Furthermore, we analysed genetic differentiation between populations of alpacas and llamas, focusing on three specific areas: northern Puno (wet Puna), southern Puno (dry Puna), and Quimsachata (the bank of germplasm in Puno). We conducted an analysis of molecular variance (AMOVA) to identify the distribution of genetic variance and the population structure, using the poppr v.2.9 package (Kamvar *et al*., 2014).

### Phylogenetics and population structure analysis

The sequences acquired were aligned with those reported in GenBank (see Supplementary S2 for guanacos, vicunas, alpacas, and llamas), utilising MAFFT v7 (Katoh *et al*., 2002). This alignment was employed to extract the haplotypes, and a maximum likelihood phylogenetic tree was constructed with 1,000 bootstrap permutations using RAXML v8.2 (Stamatakis, 2014). A principal components analysis (PCA) was conducted to examine population differentiation among the haplotypes of alpacas, llamas, and their ancestors. To assess the maternal contribution of the wild ancestors (guanaco and vicuna) in the studied populations, a Median Joining Network was performed using fastN v.1.2.4 (Chi *et al*., 2023). Lastly, a Sankey diagram was performed using the Plotly library in Python (Plotly Technologies Inc., 2015) to visualise the flow of maternal haplotypes between species and breeds.

## RESULTS

### Studied populations showed high genetic diversity within species and across agroecologies

A total of 688 sequences were analysed for domestic camelids, leading to the identification of 41 variation sites within a 468 bp alignment (Supplementary Table S2). A total of 55 haplotypes were defined across both species. Among these, alpacas and llamas shared 38% of the haplotypes present in the region, with H004 being the most common, comprising 103 individuals (40 llamas and 63 alpacas). Additionally, haplotypes H002 and H005 were the second most frequent (Supplementary Table S3).

Among the various agroecologies, the dry puna has the highest number of haplotypes, totalling 38. The wet puna follows, with 32 haplotypes, while the Quimsachata germplasm bank contains only 19 haplotypes (see Figure 2a). Notably, the dry puna also has the largest number of unique haplotypes—those not shared with other groups—amounting to 19, compared to 12 unique haplotypes in the wet puna and just 4 in Quimsachata. This distribution indicates that, although haplotype diversity is greater in the wet puna overall, the dry puna maintains a higher count of unique, non-shared haplotypes. Furthermore, Quimsachata hosts approximately 42% of the haplotype diversity found across the entire region.

**Figure 2.**
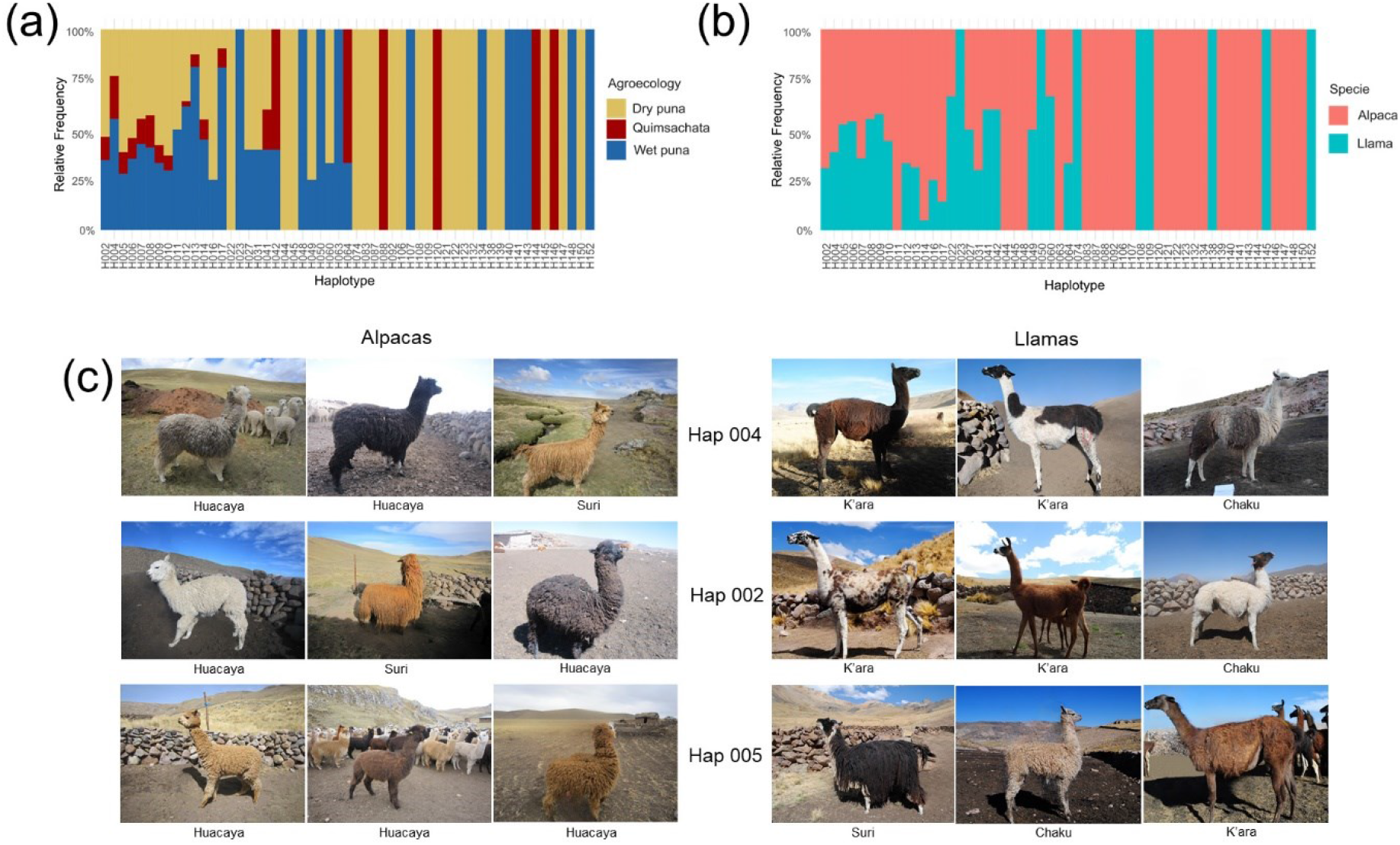
(a) Haplotype distribution across studies in agroecological zones. (b) The haplotype distribution found in alpacas and llamas was analysed. (c) Phenotypic diversity in llamas and alpacas containing the most frequent maternal haplotypes in the Southern region of Peru. (The authors captured photographs shown during sampling.)

When comparing the identified haplotypes among species (see Figure 2b and Table 1), the alpaca population exhibited a total of 47 haplotypes, 26 of which are specific to alpacas and not shared with llamas. In contrast, the llama population comprised a total of 21 haplotypes, including eight that are unique to llamas (see Supplementary Table S3). Figure 2(c) illustrates the phenotypic diversity among different breeds of alpacas (Huacaya and Suri) and llamas (Chaku, K’ara, and Suri), showcasing the most prevalent haplotypes, which are present in 39% of observed llamas and 38% of observed alpacas.

**Table 1.** Genetic diversity parameters among the tested populations.

| Specie | Population | Sample Size | Variable sites | Number of Haplotypes | Haplotype diversity ($h$) | Nucleotide diversity ($\pi$) | $Ti/Tv$ ratio |
| --- | --- | --- | --- | --- | --- | --- | --- |
| *Vicugna pacos* | Wet puna | 186 | 32 | 2518 | 0.915 | 0.019 | 7.25 |
|  | Dry puna | 175 | 35 | 32 | 0.927 | 0.018 | 8 |
|  | Quimsachata | 45 | 28 | 18 | 0.912 | 0.019 | 6.25 |
|  | Total | 406 | 41 | 47 | 0.925 | 0.019 | 9.5 |
| *Lama glama* | Wet puna | 104 | 31 | 20 | 0.912 | 0.015 | 7 |
|  | Dry puna | 139 | 35 | 22 | 0.894 | 0.017 | 8 |
|  | Quimsachata | 39 | 24 | 8 | 0.835 | 0.016 | 5.25 |
|  | Total | 282 | 37 | 29 | 0.908 | 0.016 | 8.5 |

Genetic diversity analyses identified 41 variation sites in alpacas and 37 in llamas (see Table 1). Both species exhibited high haplotype diversity across various agroecologies and within the germplasm bank (*h* ≥ 0.9). However, llamas from the dry puna and Quimsachata regions displayed slightly lower haplotype diversity (*h* ≈ 0.8). Additionally, nucleotide diversity was marginally lower in llama populations (*π* = 0.016) compared to alpaca populations (*π* = 0.019), indicating that alpacas possess a higher level of genetic diversity than llamas.

The transition-to-transversion ratio (*Ti/Tv*) obtained (Table 1) suggests that this region is more susceptible to transition mutations than transversions in both alpacas (*Ti/Tv* = 9.5) and llamas (*Ti/Tv* = 8.5). However, populations of llamas and alpacas from the Quimsachata germplasm bank display a lower ratio (5.25 and 6.25) compared with their dry and wet puna counterparts.

### The population of alpacas and llamas does not show recent demographic changes

We conducted two neutrality tests and a mismatch distribution analysis on the data to examine the potential effects of evolutionary forces on population demographics over time. To compare the observed nucleotide diversity with the expected value under the assumption of neutrality, we calculated the Tajima test (Tajima, 1989), which yielded positive values for both alpacas (*Tajima* = 1.140, *p-value* = 0.25) and llamas (*Tajima* = 0.618, *p-value* = 0.52). However, the p-values were not significant. Next, we employed the Ramos-Onsins-Rozas test *R*^*2*^ (Ramos-Onsins & Rozas, 2002), which assesses how well the data aligns with neutral expectations. In this case, both llamas and alpacas exhibited low values; however, significance was not observed, with R2 values of 0.098 (p-value = 0.73) and 0.106 (*p-value* = 0.83), respectively. In addition, the histogram of pairwise differences between sequences illustrates the mismatch distribution for both llama and alpaca populations (see Supplementary Figure 1). For both species, a multimodal distribution is evident, characterised by multiple peaks, which indicates that these populations do not exhibit evidence of recent or large-scale demographic change.

### Non-population structure was displayed within alpacas and llamas from Southern Peru

Analyses on the studied populations indicate low genetic differentiation between alpacas and llamas. To assess this, we applied the fixation index (*Fst*) to quantify genetic differentiation between populations and Nei’s genetic distance to measure divergence. Overall, alpaca and llama populations exhibited little genetic differentiation, with *Fst* values ranging from 0.01 to 0.05 (see Supplementary Table S4). Interestingly, when comparing alpacas and llamas from dry and wet puna, a similar genetic differentiation was observed (*Fst* = 0.02 – 0.03). Conversely, the Quimsachata germplasm bank exhibited similar genetic differentiation compared to wet puna for both alpacas and llamas (*Fst* = 0.01). However, it demonstrated higher genetic differentiation with llamas from dry puna (*Fst* = 0.05) than with alpacas from wet puna (*Fst* = 0.02). *Nei’s* genetic distance supports the same patterns of divergence between both species and agroecologies (Supplementary Table S4), confirming that the most significant differences are observed between llamas from dry puna and individuals from the Quimsachata bank.

To understand the genetic variation within and among the different studied populations, analysis of molecular variance (AMOVA) was performed. This analysis revealed no population structure among the Dry Puna, Wet Puna, and Quimsachata Bank populations for both alpacas and llamas. For alpacas and also llamas, variation within samples explained most of the variation, with 98% of variation for alpacas and 92% for llamas, respectively, although the p-values were not significant. (See Supplementary Table S5).

Additionally, a principal component analysis (PCA) was implemented to visualise the variation among the data. Figure 3(a) illustrates PC1 and PC2, which explain 74.6% of the variation among the studied individuals. Two main clusters were identified. Interestingly, the agroecology component does not display any effect on the structure of the PCA, as both clusters (1 and 2) contain individuals from both species and from dry and wet puna, as well as from the Quimsachata bank indistinctively.

**Figure 3.**
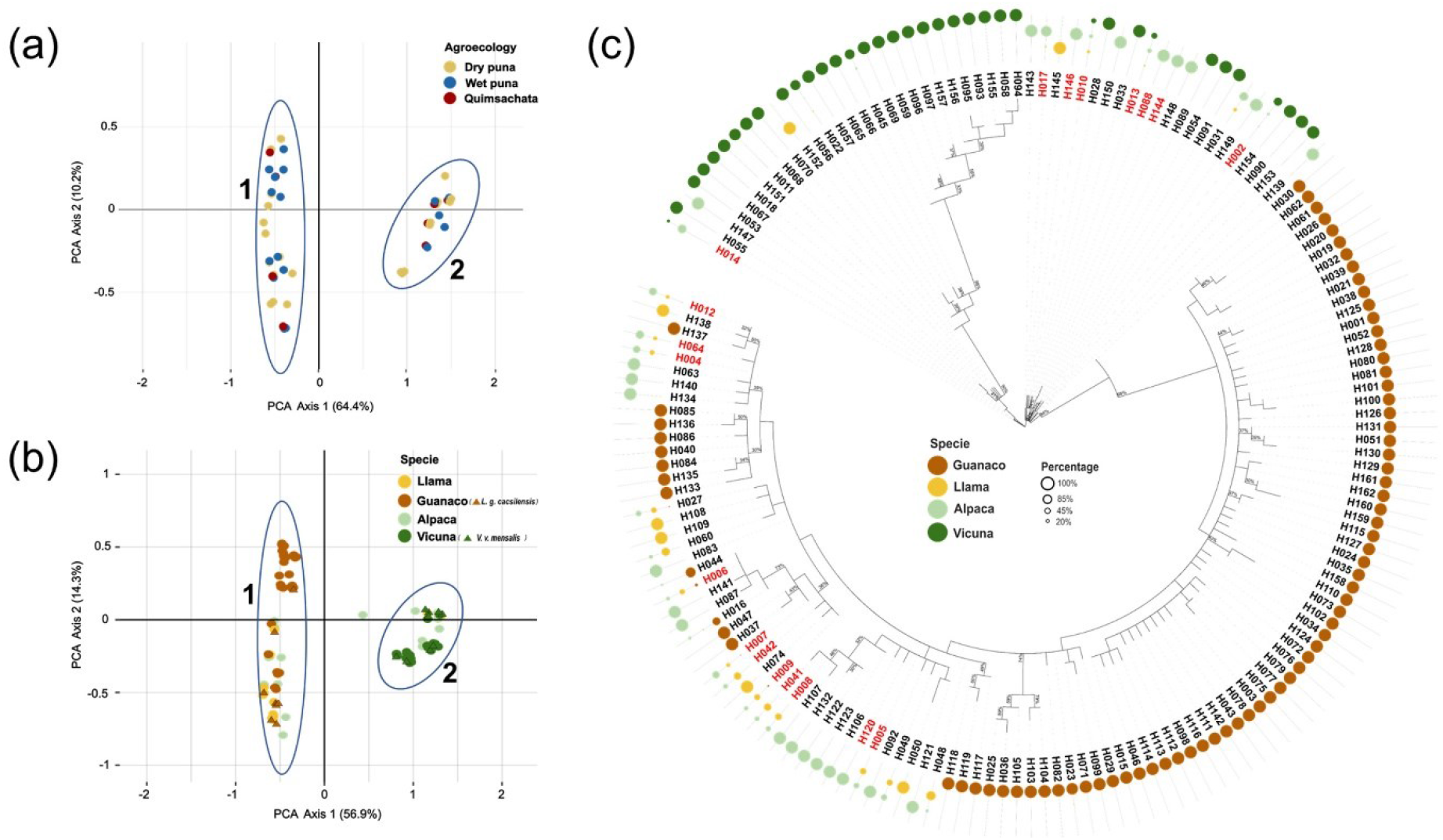
a) Principal component analysis for the studied populations of llamas and alpacas classified into dry puna, wet puna, and the Quimsachata germplasm bank. b) Principal component analysis among the studied samples (classified by species) and available sequences of South American camelids from the NCBI database. c) A phylogenetic analysis of detected haplotypes across species of South American camelids, with circles representing the frequency of each haplotype by species, and red labels indicating haplotypes observed in Quimsachata.

### Phylogeny against other domestic and wild South American camels confirms interspecific hybridisation

To understand the extent of mitochondrial variation present in llamas and alpacas from the southern region of Peru and the Quimsachata bank, phylogenetic analyses were conducted. This aims to scrutinise the haplotype diversity exhibited by the studied populations, particularly within the germplasm bank. Given that the wild ancestors of alpacas and llamas, vicunas and guanacos, share a significant autosomal and mitochondrial genetic component with their domestic counterparts, these species were also considered.

A total of 1,852 sequences from South American camelids (domestic and wild) were aligned (see Supplementary Table S6). This resulted in a final alignment of 301 bp, leading to the identification of a total of 162 distinct haplotypes (Supplementary Tables S7, S8). Unique sequences for each haplotype were considered for further analyses.

A PCA of the entire set of haplotypes was conducted (Figure 3b). PC1 and PC2 accounted for 71.2% of the variation. Interestingly, the same structural pattern observed previously in Figure 3(a) (clusters 1 and 2) was also identified when considering the total haplotypes, including wild and domestic sequences from other databases. In this analysis, cluster 1 comprises haplotypes from llamas and guanacos, consistent with their evolutionary ancestry. Notably, several alpaca haplotypes were also included in this cluster. Conversely, cluster 2 contains haplotypes from alpacas and vicunas, which aligns with their evolutionary lineage, and it also includes a small number of llama haplotypes.

The topology of the phylogenetic tree of these haplotypes confirms the same two clusters illustrated in the PCA (see Figure 3c), with the guanaco-llama branch being closely related to 19 haplotypes identified in alpacas. Similarly, the vicuna-alpaca branch retains eight haplotypes identified in llamas, confirming maternal hybridisation in both directions, from guanaco-llama to alpacas and vicuna-alpaca to llamas.

The phylogenetic relationships indicate that the most abundant haplotypes, referred to as H001, do not originate from a common evolutionary ancestor. Specifically, H001 is found in guanacos, whereas H002 is shared among llamas, alpacas, and vicunas. The haplotype identified as H003 is present only in guanacos, and H004 is exclusive to domestic camelids. In contrast, the haplotypes H017, H028, H064, H049, H041, H042, H060, and H048 are specific to domestic camelids but are found in lower abundance, with no evidence indicating unique evolutionary origins.

### Quimsachata germplasm bank preserves the most abundant haplotypes within the region

To visualise the genetic relationships among the various haplotypes identified, a median-joining network is presented in Figure 4(a). It provides clear relations within haplotypes that illustrate part of the evolutionary history of these haplotypes and their connections within cluster 1 (now defined as the guanaco haplogroup) and cluster 2 (now defined as the vicuna haplogroup), based on the species’ ancestry.

**Figure 4.**
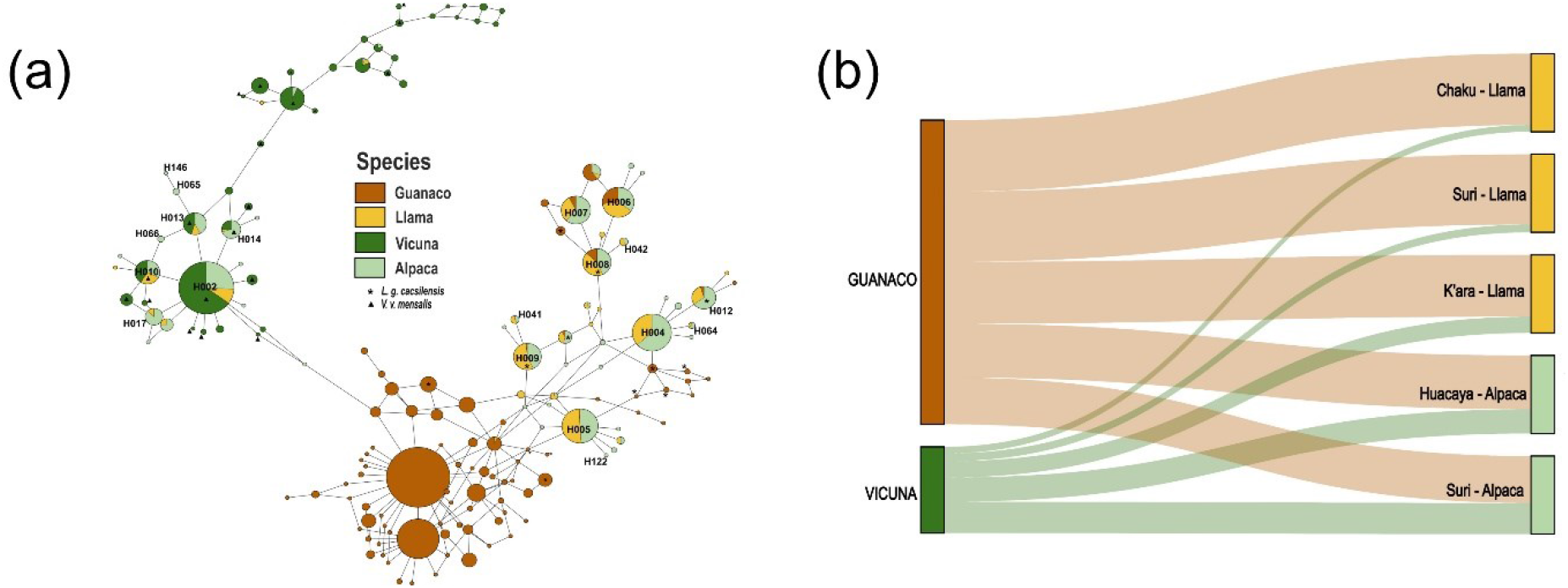
(a) Median Joining Network of alpacas, llamas, vicunas and guanacos using the D-loop region of mtDNA. The (*) and (Δ) symbols represent haplotypes with sequences from different subspecies. Haplotype IDs are included only if they contain sequences from GenBank Quimsachata. (b) A Sankey plot represents the mitochondrial haplotype flow between the different South American camel samples analysed in this study, considering species and breed within each species.

The network indicates that the guanaco group possesses a greater number of haplotypes and more complex interactions compared to the vicuna group. Within the vicuna haplogroup, several of the most common haplotypes (H002, H010, H013, and H014) include most alpaca haplotypes. Conversely, in the guanaco haplogroup, numerous llama haplotypes, such as H004 and H005, are absent from the most frequent haplotypes.

The network analyses revealed no distinct structure within the dry and wet puna or the Quimsachata germplasm bank. Instead, the analyses supported the distribution of ancestral maternal haplogroups across the samples. Furthermore, it demonstrated that the most prevalent haplotypes identified within these haplogroups are also represented by individuals conserved in the Quimsachata germplasm bank (Supplementary Table S1).

A Sankey plot of the analysed southern Peruvian samples examined here (Figure 4b) further supports this hybridisation between species, indicating a greater incidence of hybridisation in alpaca individuals carrying a guanaco haplotype compared to llamas possessing vicuna haplotypes. Among llama breeds, Chaku, K’ara, and Suri individuals exhibited the least contribution of alpaca maternal heritage, with 8.5%, 20.7%, and 10%, respectively, in contrast to the Huacaya and Suri alpaca breeds, which demonstrate 69% and 60% of maternal cross-hybridisation, respectively.

## DISCUSSION

In the southern Peruvian highlands, domestic camelids serve as a vital source of labour and income for local farmers. Traditionally, medium and small-scale alpaca and llama farmers have relied on phenotypic assessment to trade animals, particularly breeding males, among themselves. This practice aims to enhance the sustainable productivity and genetic quality of their herds. It is characterised by: (i) the use of high genetic quality males, (ii) an animal loan and sales system, (iii) the exchange of breeding stock based on trust and interpersonal relationships between communities, and (iv) the promotion of genetic improvement (FAO, 2005).

In the southern region of Peru, farmers’ associations supervise and guide agricultural practices. However, it is essential to go beyond phenotypic assessments to ensure that genetic improvement is achieved and genetic drift does not occur. This study analyses one component of this diversity by evaluating maternal haplotype diversity through the analysis of the D-loop mtDNA in alpacas and llamas from the southern region of Peru. The goal was to understand how breeding and trading systems could affect their genetic diversity and to determine whether sufficient genetic variation is maintained across herds. Additionally, we focused our analyses on the alpaca and llama herds housed in the Quimsachata ex situ germplasm bank to compare their diversity and verify the extent to which this diversity is being preserved.

Mitochondrial DNA variation in SAC has been, and continues to be, a focal point for assessing genetic relationships, studying population diversity, and addressing phylogenetic and evolutionary questions (Wheeler *et al*., 1995, 2004; Mate *et al*., 2004; Marin *et al*., 2007, 2008; Barreta *et al*., 2013; Cassey *et al*., 2018; and Abbona *et al*., 2020). Consequently, the domestication process and the relationships between species and their wild ancestors have already been explored.

In our study, alpacas and llamas displayed high levels of haplotype and nucleotide diversity. This was observed in herds from the ex situ bank and those raised by local farmers in the region. Barreta *et al*. (2013) found similar results for alpacas and llamas in Bolivia, which is geographically close to southern Peru. Additionally, Díaz-Moroto *et al*. (2021) reported similarly high values when comparing modern and ancient samples. These findings suggest that a significant maternal lineage diversity has been preserved throughout the region since domestication, despite the various evolutionary and population structure events that have impacted these species since the arrival of the Spanish conquerors (Renieri *et al*., 2009; Díaz-Maroto *et al*., 2021).

Moreover, our AMOVA analyses revealed no evidence of population structure, indicating that most genetic variation exists within individuals rather than between them. This suggests that individuals are likely mixing freely across groups (or that the farmer’s practices maintain a constant movement between herds), resulting in high genetic flow, low intrapopulation differentiation, and interconnected populations. This pattern corresponds with the breeding practices of farmers in the area, who trade animals.

South American camels have undergone various demographic changes throughout different periods. For example, bottlenecks have been estimated for wild SAC after the mid-Holocene (Casey *et al*., 2018). Similarly, following the arrival of Europeans on the continent, the SAC population was significantly reduced in both number and geographic distribution (Orlove, 1977; Díaz-Lameiro *et al*., 2022). Flores Ochoa (1982) estimated that approximately 90% of the SAC were diminished after the Spanish conquest. Conversely, our analyses to assess whether any strong demographic event has occurred over the sampled populations resulted in values that might suggest a past demographic change; however, we found no statistical significance. This is likely attributable to the nature of population dynamics, such as introgression events during pre-conquest domestication (Díaz-Lameiro *et al*., 2022) and post-conquest interbreeding (Wheeler *et al*., 1995); as well as to the modern trading and breeding systems maintained by local farmers (FAO, 2005).

Mitochondrial variation not only confirms the domestication origin of llamas and alpacas (Marin *et al*., 2007, 2017) but also indicates interbreeding among species. Using microsatellite markers, Varas *et al*. (2020) have provided evidence of this interbreeding, while Echalar & Barreta (2022) have measured not only the introgression between species but also the generation and maintenance of fertile hybrids known as “huarizos” within the herds. Our results display the flow of maternal haplotypes among species, with over half of the maternal alpaca hybrids, in contrast to around 10 to 20% of the llama hybrids. This finding is consistent with those of Marin *et al*. (2017), Varas *et al*. (2020) and Pallotti *et al*. (2023), suggesting that recent and more common interbreeding has occurred between alpaca males and llama females than vice versa.

*Ex-situ* or off-site conservation complements in situ efforts as a backup strategy for maintaining genetic diversity, alongside other activities such as recovery, reintroduction, and research (McGowan *et al*., 2017). Since its establishment in 1988, the Quimsachata germplasm bank has served as a focal point not only for preserving coloured alpacas (non-white) but also for developing shearing, breeding, and reproductive technologies beneficial to farming communities in the region (Huanca *et al*., 2007). Significant concerns regarding this practice, aside from maintaining the facilities and staff, as well as other associated costs, include the potential for genetic decline and reduction of effective population size due to selective breeding and limited space (Frankham *et al*., 2010). Genetic diversity analyses using microsatellites on llamas (Paredes *et al*., 2020) and alpacas (Figueroa *et al*., 2023), including individuals from the Quimsachata bank, have revealed high levels of diversity, even greater than those in herds maintained by local farmers in the Southern Peruvian region.

On the mitochondrial diversity side, our study has also revealed that the Quimsachata bank currently holds a significant maternal heritage diversity of alpacas and llamas in the region, even though the sample size is relatively smaller compared to the herds of alpacas and llamas analysed here. These findings provide important evidence of the bank’s positive efforts in preserving this diversity and mitigating the potential effects of genetic drift, selective breeding or low migration rates.

Our study also considered agroecology (wet and dry puna) as a potential factor contributing to constraints on the preservation of genetic diversity. The results indicated that Quimsachata maintains slightly less genetic distance with herds in the wet puna than with those in the dry puna. Furthermore, dry puna llama and alpaca herds contain a greater number of private haplotypes specific to this agroecology, which were not found in either the wet puna herds or in Quimsachata. In comparison with wet puna, and despite rural development programmes, poverty remains a significant challenge for the alpaca and llama communities of dry puna in southern Peru. This situation entails reduced access routes and organisational farm systems, which could impact the trading band breeding systems of South American camels (Leyton *et al*., 2019). The maternal private diversity observed in this agroecology may reflect farmers’ limited resources for exchanging breeding animals and promoting genetic improvement.

Although our study only encompasses the maternal diversity and interbreeding assessment in llamas and alpacas from Southern Peru, it fully addresses how this diversity is maintained across the region, which has the highest production of alpacas and llamas worldwide. Additionally, it contrasts this with the ex situ conservation efforts led by the Ministry of Agriculture in Peru. These results are significant for camelid breeders and industries globally, given the economic importance of their products. By providing these findings, farmers, their families, and livelihoods can benefit, as this information can contribute to breeding strategies, which is particularly crucial due to the often-impoverished Andean rural communities, especially those living in the dry puna, including the South Puno, Moquegua, and Tacna regions. Further efforts are still needed in this area, and further genomic approaches like whole genome sequencing (WGS), genomic structural variation (SV), and genetic expression studies, such as RNA sequencing and methylation analyses, will enable a refined understanding of this diversity and assist in developing a more informed management of genetic diversity across species and selection programmes.

## CONCLUSION

Our data indicate a high level of genetic diversity and low genetic differentiation in alpaca and llama populations from the southern region of Peru. This diversity remains significant even when compared to the maternal haplotypes of domestic and wild SAC found in other regions of Peru and South America, where these animals are typically distributed. No recent demographic changes or alterations in population structure were observed within the analysed populations. Furthermore, our data confirm two specific clusters among alpacas and llamas, reflecting their ancestral relationships with the haplogroups of vicunas and guanacos. However, this distribution also suggests a notable degree of crossbreeding between the two species, with alpacas exhibiting a more substantial amount of maternal ancestry from guanaco-llamas. Preserving the diversity of South American camels is a key component for the economic development of Andean communities. Thus, the ex situ efforts assessed here have shown a positive impact on maintaining this diversity. Further work on in situ and ex situ herds will be necessary to dissect the complex diversity of these species, but more importantly, to study vital productivity traits that can influence the sustainable productivity of these animals, thereby ensuring greater benefits for farmers while also preserving the genetic variation of individual animals.

## Data availability statement

The resulting sequences used in this study are publicly available via the project number 2939020 in GenBank (NCBI) with accession numbers PV359989-PV360677.

## Authorship contribution

**O.M**. sampling, wet lab, data analysis, data visualisation, writing and editing. **D.C**. Sampling, wet lab, data analysis. **C.Y**. Wet lab, review and editing. **T.H**. Sampling supervision. **N.A**. Sampling, sampling supervision. **E.S**. Sampling, review and editing. **A.V-T**. Conceptualisation, project administration, sampling, data analysis, and led the writing of the manuscript, although all authors contributed critically to the drafts.

## Acknowledgments

The authors would like to thank all the alpaca and llama breeders who contributed to this work by facilitating coordination and access to the samples. Special thanks to Mr Maximo Odon Díaz Bustinza for his extensive support, including logistics and access to his herds. Additionally, we extend our gratitude to all the staff at the Quimsachata station, whose invaluable efforts were crucial to the completion of this work. We wish to dedicate this work to our friends and colleagues Diogenes Cerna, Dr Teodosio Huanca, and Ing. Norberto Apaza, who contributed immensely to this project, and whose memory will be cherished within the South American Camels scientific community, as well as by all those who had the privilege of knowing them.

