## Supplementary Figure for "Assessment of Ex Situ Conservation Strategies for the Preservation of Domestic South American Camels in Southern Peru: Insights from Mitochondrial DNA Analysis"

### Supplementary Figures

A) B)

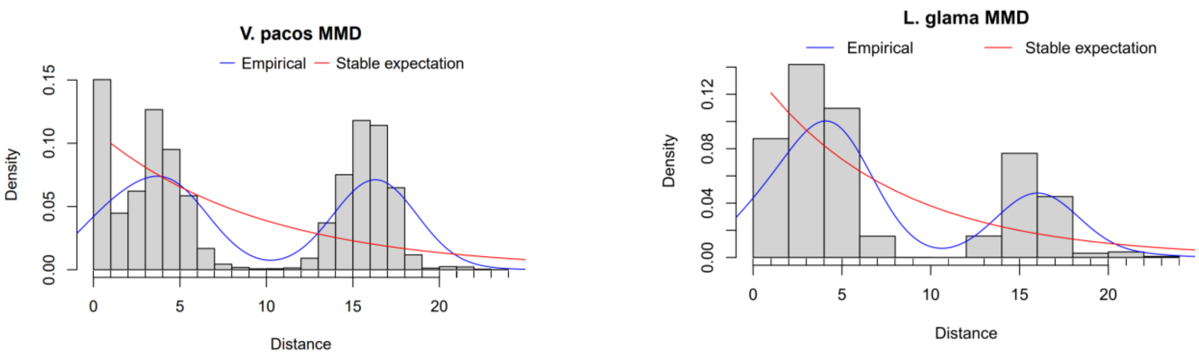


**Figure S1**. Total mismatch distribution of a) Alpacas, and b) Llama populations of the Southern Peruvian region.
